# Modeling nonsegmented negative-strand RNA virus (NNSV) transcription with ejective polymerase collisions and biased diffusion

**DOI:** 10.1101/2022.06.24.497531

**Authors:** Felipe-Andrés Piedra

## Abstract

Infections by nonsegmented negative-strand RNA viruses (NNSV) are widely thought to entail gradient gene expression from the well-established existence of a single promoter at the 3’ end of the viral genome and the assumption of constant transcriptional attenuation between genes. But multiple recent studies show viral mRNA levels in infections by respiratory syncytial virus (RSV), a major human pathogen and member of NNSV, that are inconsistent with a simple gradient. Here we integrate known and newly predicted phenomena into a biophysically reasonable model of NNSV transcription. Our model succeeds in capturing published observations of RSV and vesicular stomatitis virus (VSV) mRNA levels. We therefore propose a novel understanding of NNSV transcription based on the possibility of ejective polymerase-polymerase collisions and, in the case of RSV, biased polymerase diffusion.

## Introduction

Viruses with nonsegmented negative-strand RNA genomes (NNSV) (all viruses of the order *Mononegavirales*) contain major pathogens such as Ebola, rabies, measles virus, respiratory syncytial virus (RSV), and vesicular stomatitis virus (VSV) – the latter is a highly studied bovine pathogen of the same family, *Rhabdoviridae*, as rabies virus.

The RNA genomes of NNSV are coated in nucleoprotein and support both whole genome replication and the transcription of subgenomic mRNAs by viral RNA-dependent RNA polymerases (pols) in the cytosol of infected cells. These genomes have a single promoter located at the 3’ end that is essential for both processes, presumably by facilitating the transient dissociation of terminal genomic RNA from nucleoprotein and the entry of viral pols, hitherto bound only to the nucleoprotein of the ribonucleoprotein (RNP) complex, into the RNA genome.

The textbook model of NNSV gene expression predicts a transcription gradient from 1) obligatory pol entry at the 3’ end of the genome; 2) start-stop transcription in response to conserved gene start (GS) and gene end (GS) signal sequences; and 3) transcriptional attenuation between genes (1, 2). However, multiple published studies show NNSV gene expression patterns – especially from RSV, which is one of its most highly studied members – that are either non-gradient, with one or more downstream genes appearing more highly expressed than upstream genes, or inconsistent with a simple gradient from a constant level of attenuation between genes (3-9). Regarding the latter, multiple studies show an abrupt and dramatic decrease in gene expression over the last two genes of the RSV genome (3-6, 9), the sole region of the genome containing overlapping ORFs – the textbook model of NNSV transcription offers no way of explaining this.

Here we implement a coarse-grained, mechanistic and stochastic computational model incorporating known and, ultimately, newly proposed features (ejective pol-pol collisions and 5’ biased pol diffusion) of the underlying molecular biophysics to gain a deeper understanding of NNSV transcription and to capture, for the first time, experimentally observed non-gradient RSV and gradient VSV gene expression patterns.

## Results and Discussion

### i. Determining the effects of stochastic transcription using a range of initiation probabilities

We took a heuristic approach to fitting actual observations of RSV and VSV gene expression and started by modeling a single pol taking an unbiased random walk down either genome and stochastically initiating and terminating transcription (Fig1A and B).

**Fig 1.**
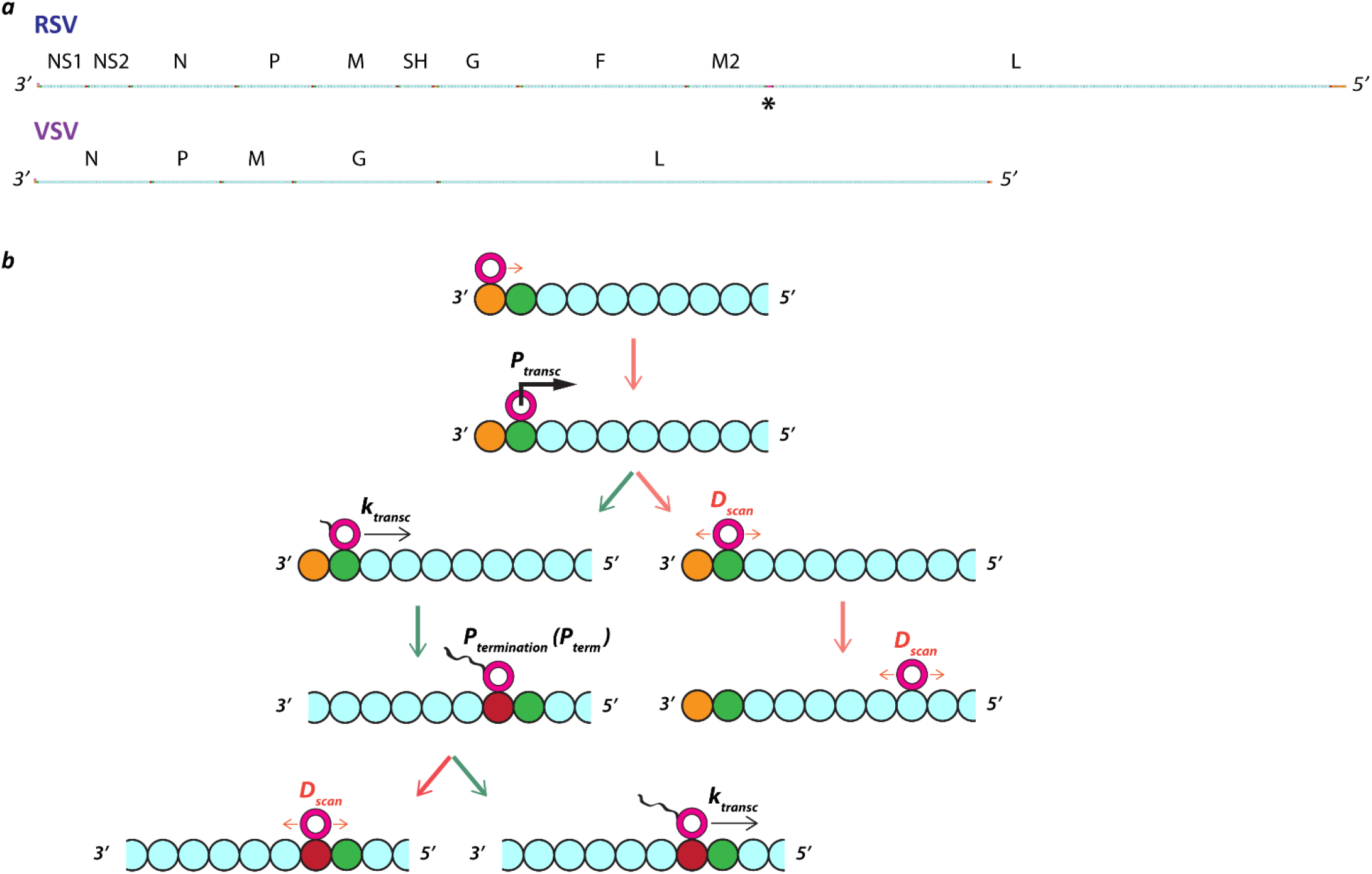
The model: linear respiratory syncytial virus (RSV) and vesicular stomatitis virus (VSV) genomes support the stochastic initiation and termination of transcription by a diffusing viral RNA-dependent RNA polymerase (pol). **(A)** The genetic structure of RSV and VSV genomes. The modeled RSV genome is 15,222 nt long and contains 10 ORFs with 8 gene junctions and a single short region (68 nt) of overlapping ORFs between genes M2 and L (see black asterisk). The modeled VSV genome is 11,152 nt long and contains 5 ORFs with 4 gene junctions. The genomes were divided into chunks approximating the size of a pol footprint (28 nts). Most of each genome is coding sequence (represented as cyan beads). **(B)** Essential model phenomena and parameters. A single RNA-dependent RNA polymerase (pol) starts an unbiased random walk at a rate *D*_*scan*_ (= 1 genomic chunk per event) at the most 3’ chunk (depicted as a burnt orange bead) of the modeled genome. Transcription initiation occurs with a probability *P*_*transc*_ when a pol diffuses onto a genomic chunk containing a gene start (GS) signal (depicted as a green bead). If transcription is not initiated, the unbiased random walk (i.e., diffusion) resumes. If transcription is initiated, the modeled pol state changes and the pol starts translocating 5’ down the genome at a rate *k*_*transc*_ (= *x* genomic chunks per event). Transcription termination occurs with a probability *P*_*term*_ when a transcribing pol translocates onto a genomic chunk containing a gene end (GE) signal (depicted as a red bead). If termination occurs, the pol state changes back to non-transcribing and resumes diffusion along the genome at a rate *D*_*scan*_; if termination does not occur, the pol ‘reads through’ the GE signal and continues transcribing into the next ORF. (Cyan beads represent coding sequence.)

In this simple case, the parameters to explore are probabilities of transcription initiation and termination. The termination probabilities can be derived directly from published sequencing data for RSV, as these are simply the complement of the published readthrough rates (9). For VSV, we made use of estimates suggesting a very high probability of termination (0.99) for the GE signals modeled here (10). In contrast with termination probabilities, probabilities of transcription initiation are completely unknown. However, the three GS signals of the RSV genome modeled here have been tested in minigenomes for their relative strength of gene expression (11). These relative strengths were used to constrain the ten transcription initiation probabilities of RSV (Table 1). The five GS signals of the VSV genome modeled here were all assumed to support an equal probability of transcription initiation.

**Table 1.**
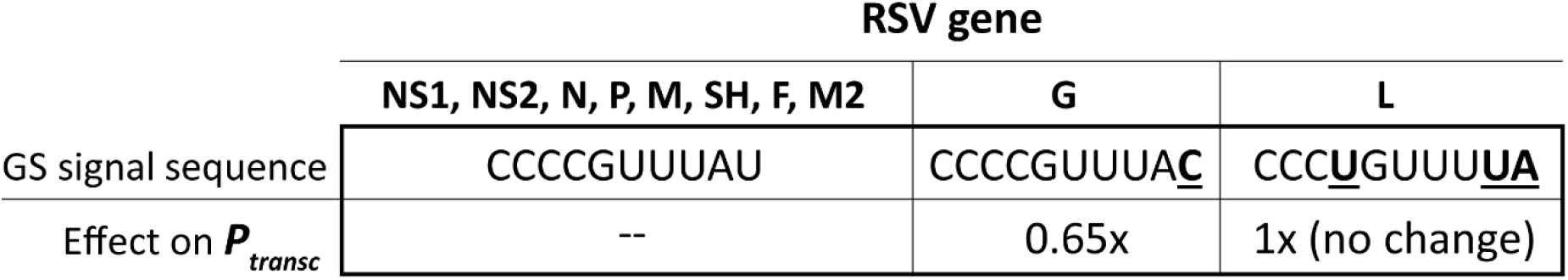
Gene start (GS) signal sequence-based constraints on RSV transcription initiation probabilities (*P*_*transc*_) from Kuo et al. Most RSV/A genomes contain three different GS signal sequences. Kuo et al. performed minigenome studies to quantify the effects on gene expression of all single nt mutations within the GS signal (11). The G gene GS signal contains a single mutation (relative to the most common GS signal sequence) at position 10 that reduced gene expression by ∼35%. Kuo et al. reported that the L gene GS signal gave rise to a magnitude of gene expression equal to that of the most common GS signal. It is therefore reasonable to model RSV transcription with a single probability of transcription initiation at all GS signals except for G, where the probability should be multiplied by 0.65.

Simulated patterns of RSV and VSV gene expression were essentially flat for all three sets of transcription probabilities (Fig2). Standard deviations of individual mRNA levels were, as expected, highest for the lowest transcription probabilities tested (Fig2). In the case of RSV, a slight bump in gene expression occurs for the SH gene and becomes most visible at the highest transcription probabilities tested (Fig2A). This is because of the lower rate of transcription initiation at the G gene GS signal (0.65x), which is directly downstream of the SH gene: the modeled pol occasionally fails to initiate transcription at the G gene before diffusing to the nearest GS signal, SH, where it is ∼1.5x more likely to initiate transcription. We also calculated a root-mean-square deviation (RMSD) for each simulated gene expression pattern to quantify how well the model fit the observed in vitro gene expression patterns (Fig 2B and D).

**Fig 2.**
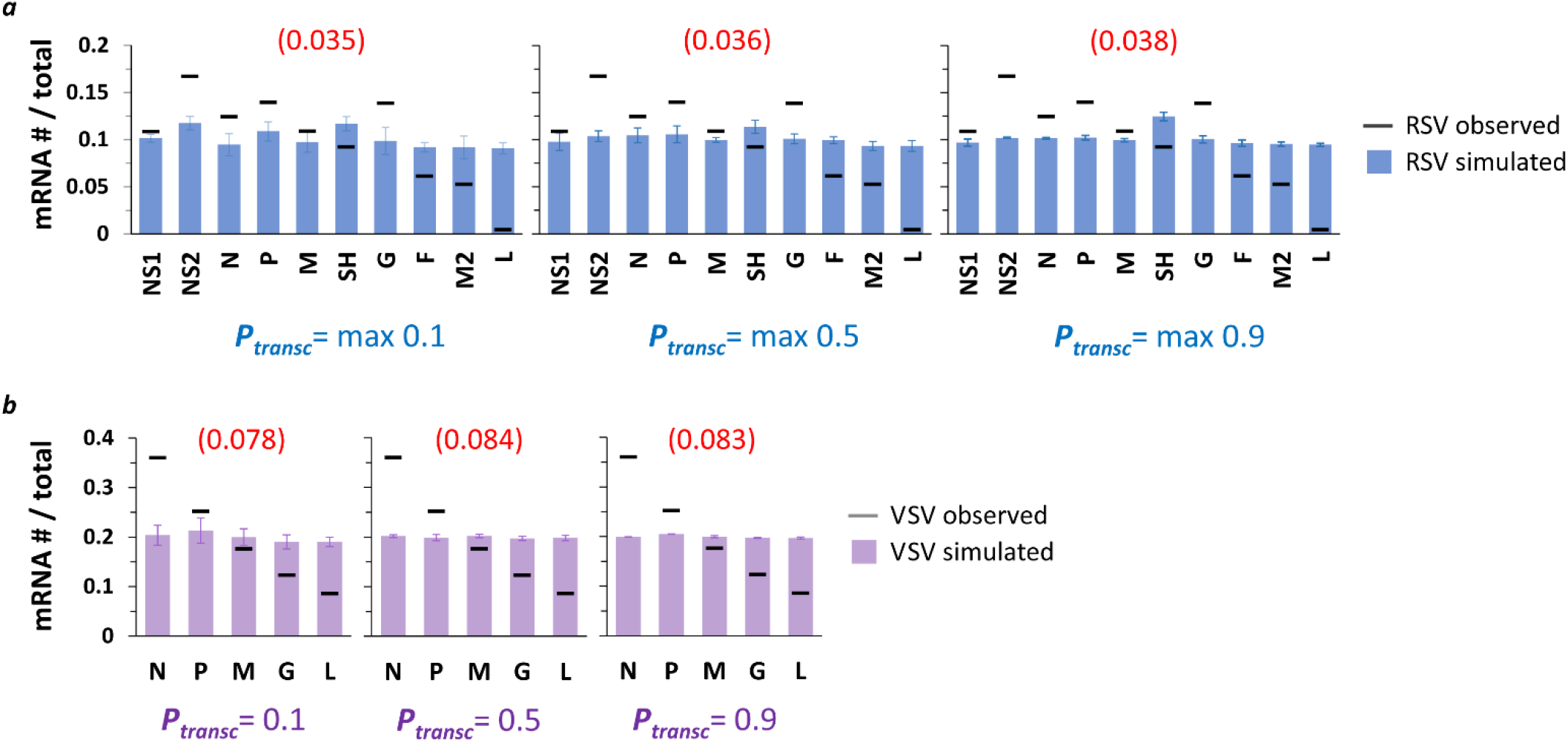
Single pol simulations produce flat patterns of gene expression across *P*_*transc*_ values tested. **(A)** Simulated RSV transcription. Histograms of mRNA # for each RSV gene divided by the total mRNA # show uniform gene expression across the 10 genes for all three sets of *P*_*transc*_ tested (max 0.1, max 0.5, and max 0.9). For each set of *P*_*transc*,_ the max value equals the probability of transcription at every GS signal except for that of the G gene, which equals 0.65*max. Blue bars depict results from simulations; black horizontal bars depict average published experimentally observed values (9). Each data point is the average of three 100,000 event simulations; error bars show the standard deviation. The number in parentheses and red above each histogram is the root-mean-square deviation (RMSD) of the simulated gene expression pattern from the experimental observations. **(B)** Simulated VSV transcription. Histograms of mRNA # for each VSV gene divided by the total mRNA # show uniform gene expression across the 5 genes for all three sets of *P*_*transc*_ tested (0.1, 0.5, and 0.9). For each set of *P*_*transc*,_ the probability of transcription is the same at every GS signal. Lavender bars depict results from simulations; black horizontal bars depict average published experimentally observed values (12). Each data point is the average of three 100,000 event simulations; error bars show the standard deviation. The number in parentheses and red above each histogram is the root-mean-square deviation (RMSD) of the simulated gene expression pattern from the experimental observations.

### ii. Incorporating multiple polymerases into our model of NNSV transcription

Modeling a single pol diffusing along an RSV or VSV genome and stochastically starting and stopping transcription with the sequence-based probabilities used here cannot capture experimentally observed gene expression patterns. It is also well established that VSV virions contain 10s of pols per genome (13), making it very likely that both VSV replication and transcription involve multiple pols interacting with a single genome.

Thus, we decided to model multiple pols interacting with and transcribing single RSV and VSV genomes. This required conceiving of rules to govern interactions between the pols interacting with a single genome. We decided to implement one-by-one pol entry at the 3’ end of the genome, a variable maximum number of pols interacting with the genome at any one time, ‘soft’ collisions between non-transcribing pols that prevent one pol from ‘hopping over’ another, and hard collisions between 5’ translocating transcribing pols and diffusing non-transcribing pols resulting in the latter’s ejection from the genome.

The latter rule was partly inspired by observing that the steepest drop in RSV gene expression, a dramatic decrease reported by multiple independent groups (3-6, 9), occurs over what should be a hot-spot for collisions between transcribing and non-transcribing pols: the overlap in the M2 and L gene ORFs (Fig3A). We also took inspiration from work by Tang et al. reporting a very high affinity of VSV pols for the VSV ribonucleoprotein (RNP) complex and suggesting, through computational modeling, the importance of a class of ejective pol-pol collisions somewhat different from the class modeled here (14). Specifically, here we model two pol states – non-transcribing, which diffuse bidirectionally; and transcribing, which move only 5’ – for pols that have gained access to the RNA genome through the 3’ promoter; in contrast, Tang et al. modeled ejective collisions between pols that have accessed the RNA genome via the 3’ promoter and pols ‘scanning’ the VSV RNP complex via interactions between pol-bound P protein and N protein for the 3’ promoter (14). We make no attempt to model the ‘scanning’ pols that have yet to access the RNA genome of Tang et al.

**Fig 3.**
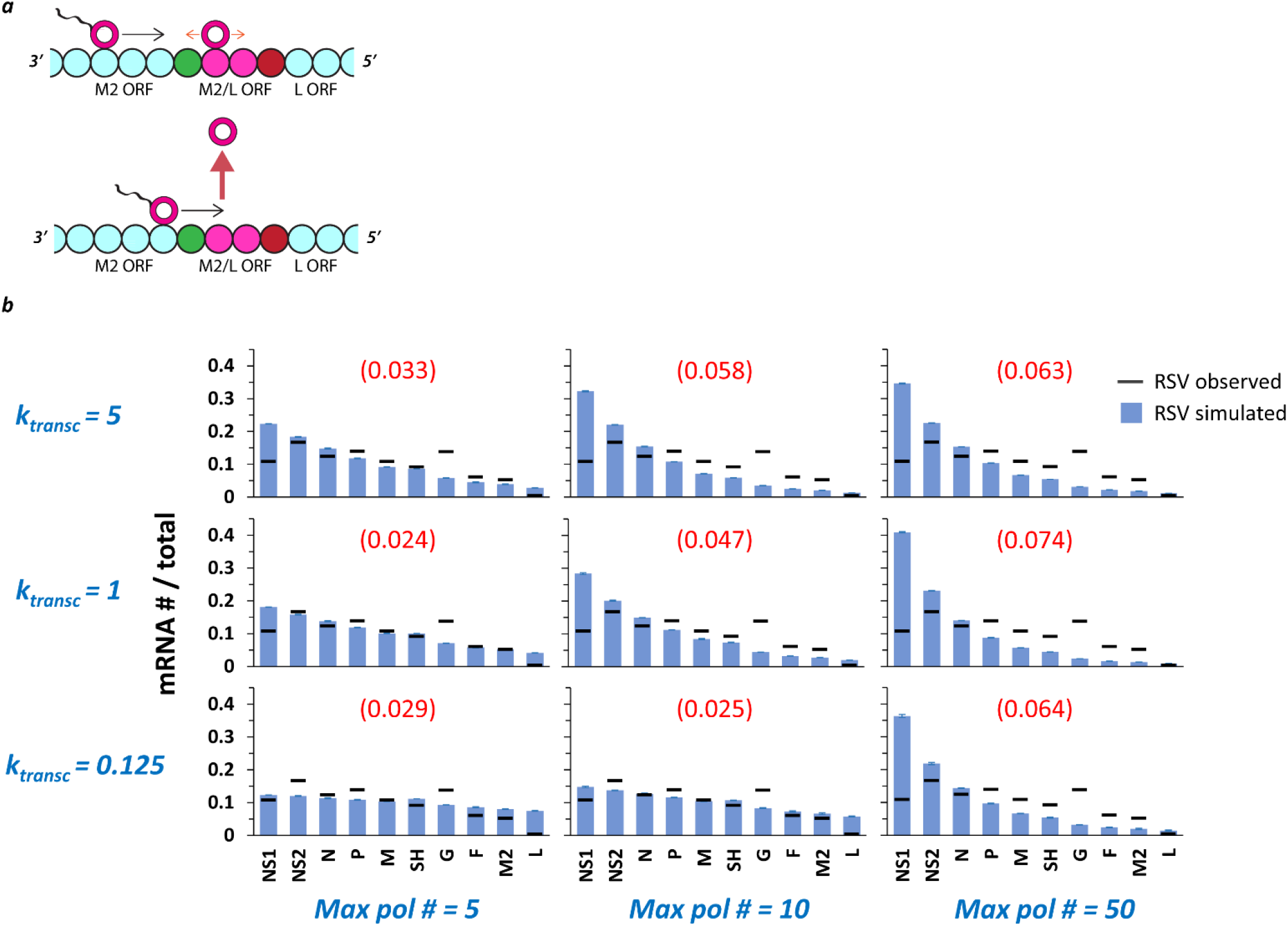
Multiple pols on a single genome undergoing ejective collisions between transcribing and non-transcribing pols produce gene expression gradients of increasing steepness with increasing 5’ translocation rate (*k*_*transc*_) and increasing *maximum pol number (max pol #)*. **(A)** The M2/L overlap in ORFs. The final two genes of the RSV genome, M2 (which encodes both a transcription processivity factor and a regulatory factor that enhances replication) and L (which encodes the polymerase), share a 68 nt stretch (approximately two genomic chunks of 28 nts each — depicted as magenta beads) of ORF. This ORF overlap should be a hotspot for collisions between transcribing pols and non-transcribing pols diffusing in the neighborhood of the M2 GE signal (shown as red bead). The L gene GS signal is depicted as a green bead. **(B)** RSV gene expression patterns over a range of *k*_*transc*_ and *max pol #*. The parameter *k*_*transc*_ sets the rate at which transcribing pols move 5’ down the genome (units = genomic chunks per simulated event) and the parameter *max pol #* sets the maximum number of pols allowed on the genome at one time. Simulations of RSV transcription were performed at three different values of *k*_*transc*_ x three different values of *max pol #*. Histograms of mRNA # for each RSV gene divided by the total mRNA # depict results from the simulations (blue bars) and average published experimentally observed values (black horizontal bars) (9). Each data point is the average of three 100,000 event simulations; error bars show the standard deviation. The number in parentheses and red above each histogram is the root-mean-square deviation (RMSD) of the simulated gene expression pattern from the experimental observations.

Because our model was modified to include multiple pols undergoing ejective collisions between transcribing and non-transcribing pols, it was necessary to explore another parameter, k_transc_, setting the 5’ translocation speed of a transcribing pol. We simulated RSV transcription under three different values each of k_transc_ and maximum pol number (Fig3B), and VSV transcription under three different values of k_transc_ and a single maximum pol number (Fig4). A single maximum pol number was used for VSV transcription because of published work suggesting approximately 50 VSV pols per VSV genome (13); to our knowledge, this ratio is not known for RSV.

**Fig 4.**
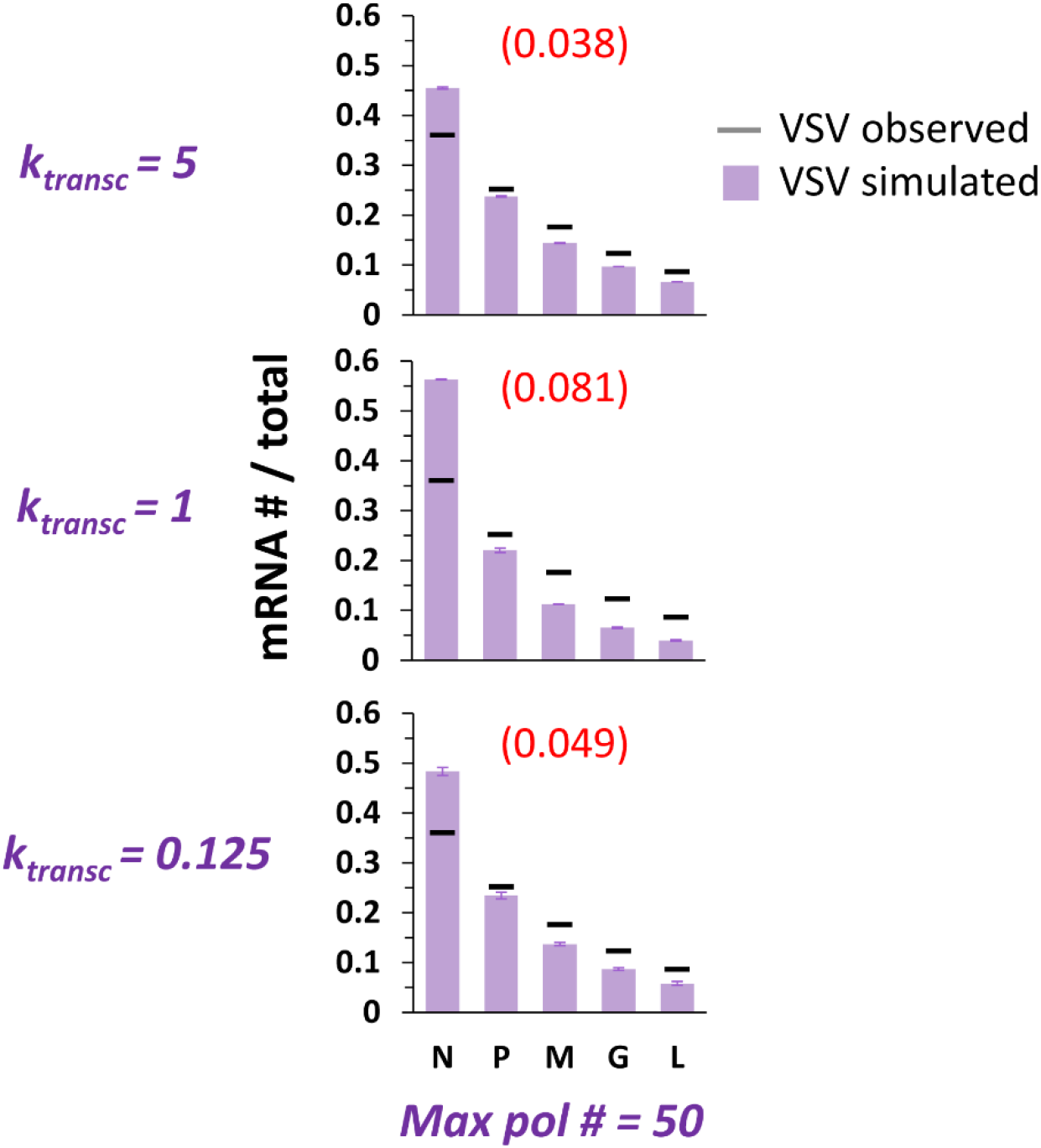
Simulations of at most 50 pols and collision-based pol ejections fit benchmark observations of VSV gene expression best at the highest *k*_*transc*_ tested. VSV gene expression patterns over a range of *k*_*transc*_ and a single *max pol #*. The parameter *k*_*transc*_ sets the rate at which transcribing pols move 5’ down the genome (units = genomic chunks per simulated event) and the parameter *max pol #* sets the maximum number of pols allowed on the genome at one time. Simulations of VSV transcription were performed at three different values of *k*_*transc*_. Histograms of mRNA # for each VSV gene divided by the total mRNA # depict results from the simulations (lavender bars) and average published experimentally observed values (black horizontal bars) (12). Each data point is the average of three 100,000 event simulations; error bars show the standard deviation. The number in parentheses and red above each histogram is the root-mean-square deviation (RMSD) of the simulated gene expression pattern from the experimental observations.

Simulated RSV gene expression patterns display a 3’ to 5’ gradient of increasing steepness with increasing maximum pol number and, for simulations with a maximum of 5 and 10 pols, with increasing k_transc_ (Fig3B). The transcription gradient in our model is a consequence of a gradient in pol concentration emerging from ejective pol-pol collisions and obligatory pol reentry at the 3’ end of the genome. In the case of simulations of at most 50 pols, the gene expression gradient is steepest at the middle value of k_transc_ because the higher value supports such a high frequency of ejective pol collisions that the actual number of modeled pols occupying a genome at steady-state tends to ∼10, while the middle value leads to one of ∼20 pols, which leads to a sharper pol concentration gradient along the genome and a steeper gene expression gradient. It is clear from both the calculated RMSD values and visually inspecting the fits that simulations incorporating a high maximum number of RSV pols per genome produce a gene expression pattern that is too steeply gradient; in contrast, simulations of at most 5 RSV pols per genome yield much better fits of the published data across the 20-fold range of k_transc_ values tested (Fig3B).

We simulated VSV gene expression across the same 20-fold range of k_transc_ values and only one value of maximum pol number (Fig4). At the highest value of k_transc_ tested, the model captures benchmark observations of VSV transcription fairly well. It is interesting that the middle value of k_transc_ results in the worst fit of the data; this results from the phenomenon described above for RSV transcription under the same maximum pol number: the highest value of k_transc_ tested leads to such a high frequency of pol collisions that the actual number of pols occupying the genome at steady-state is much lower than the maximum possible; because the lower value of k_transc_ leads to less frequent collisions and a concomitant increase in the number of pols occupying the genome, a steeper gene expression gradient results (Fig4).

### iii. Further exploring the effects of collision-based pol ejections on RSV transcription

Thus, our simple model incorporating multiple pols undergoing random diffusion along the genome when not transcribing and ejective collisions when a transcribing and non-transcribing pol meet captures benchmark observations of VSV gene expression (12) well while poorly fitting our published observations of RSV gene expression (9). Furthermore, the model most poorly fits data coming from the last two genes of the RSV genome, where multiple groups report a dramatic decrease in gene expression. This is the sole region of the modeled genomes where two ORFs overlap; and this overlap helped inspire the addition of ejective pol collisions into our model. We decided to further investigate the effect of the modeled pol collisions on gene expression over the M2-L region of RSV by analyzing the relationships between 1) the number of pol ejections per run of our simulation and values of the maximum pol number and k_transc_; and 2) ratios of L and M2 mRNA levels and values of the maximum pol number and k_transc_ (Fig5).

**Fig 5.**
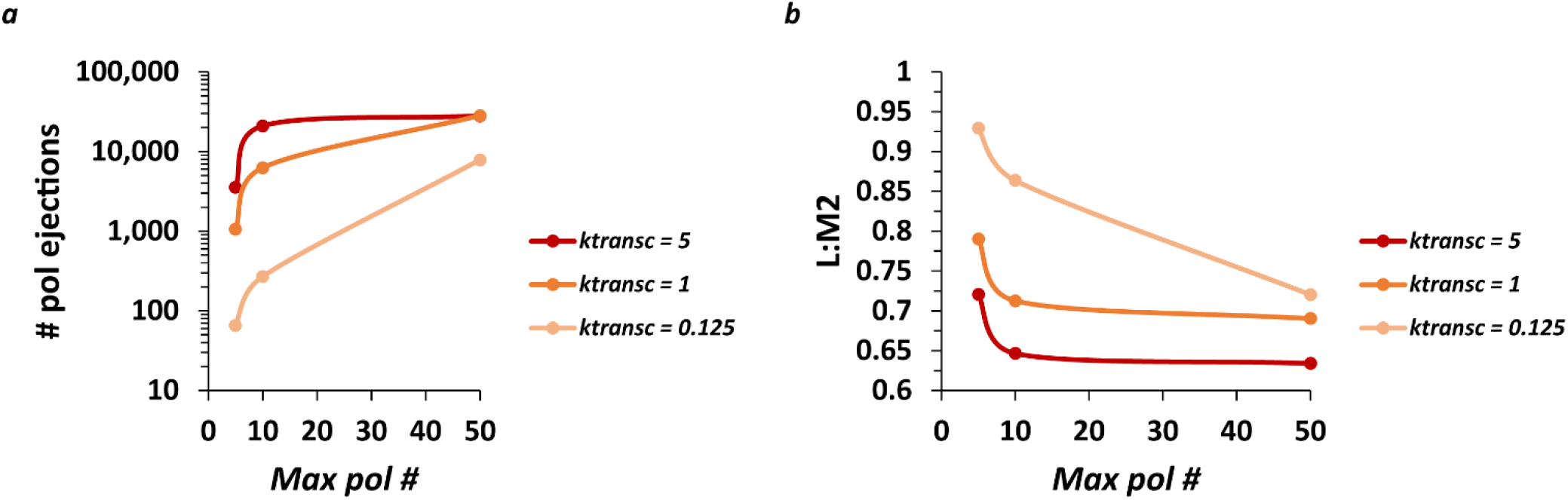
The number of pol ejections occurring over one run of the model and the ratio of RSV L mRNA to M2 mRNA levels (L:M2) produced are inversely related. **(A)** The number of pol ejections vs. *max pol #* for three different values of *k*_*transc*_. Each data point is the average of three 100,000 event simulations. **(B)** The ratio of L mRNA to M2 mRNA levels (L:M2) vs. *max pol #* for three different values of *k*_*transc*_. Each data point is the average of three 100,000 event simulations.

The average number of pol ejections generally increases with increasing max pol # and increasing k_transc_ (Fig5A). However, at the higher values of k_transc_ (1 and 5 genomic chunks per event), the average number of pol ejections starts to plateau beyond a max pol # of 10. This again shows that the steady-state number of pols bound to a genome in the model depends on k_transc_ and that this number is close to 10 at the highest value of k_transc_ tested (assuming a pol footprint of 28 nt). In addition, curves for the higher values of k_transc_ start to converge, suggesting that the model is reaching its maximum pol ejection frequency.

Consistent with the expected effect of the modeled pol collisions, ratios of L:M2 generally decrease with increasing max pol # and increasing k_transc_ (Fig5B). However, L:M2 plateaus sharply for the higher values of k_transc_ tested beyond a max pol # of 10. In addition, curves for higher values of k_transc_ start to converge, suggesting that the model is reaching its minimum L:M2 which remains much higher than the average experimentally observed value of ∼0.09 (9). This suggested that the RSV model was missing something of fundamental importance.

### iv. Modifying the model to include 5’ biased diffusion of non-transcribing pols

We therefore decided to test the effects of including biased pol diffusion in our model, specifically a 5’ bias, which might help explain why gene expression is possible but falls off steeply when a pol must diffuse 3’ to reach the nearest GS signal after terminating transcription (as occurs in RSV from the M2-L overlap) (Fig6A). The biased diffusion of proteins has been shown before (15-17), making this change to the model biophysically reasonable.

**Fig 6.**
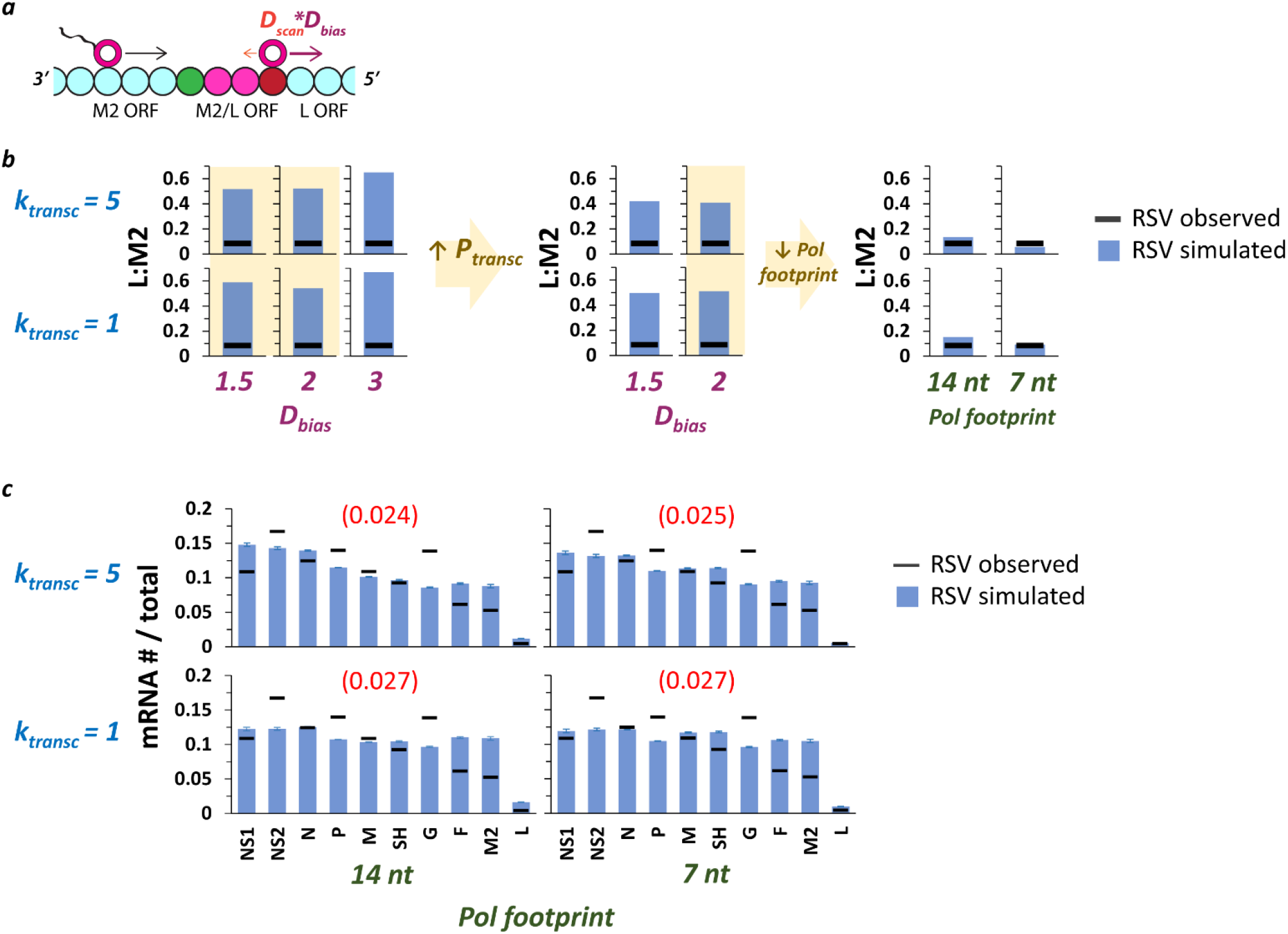
Including a 5’ pol diffusion bias (*D*_*bias*_) can reproduce the experimentally observed drop in gene expression between RSV genes M2 and L. **(A)** *D*_*bias*_ is expected to have its largest effect on ratios of L mRNA to M2 mRNA levels (L:M2). M2 and L ORFs are depicted as cyan beads (bead size reflects a pol footprint size of 28 nt); the M2/L overlap is depicted as two magenta beads; the L GS signal is depicted as a green bead; the M2 GE signal is depicted as a red bead. **(B)** A scan of parameter values shows that moderate *D*_*bias*_ with reduced pol footprint size can reproduce the experimentally observed value of L:M2 (9). Left panel: histograms show simulated (blue bar) and experimentally observed (horizontal black bar) L:M2 values for two different values of *k*_*transc*_ x three different values of *D*_*bias*_ (*max pol #* = 5). The two lower values of *D*_*bias*_ tested (highlighted in pale yellow) result in a slightly greater drop in L:M2 than *D*_*bias*_ = 3; these values were used in subsequent simulations. Middle panel: histograms show simulated (blue bar) and experimentally observed (horizontal black bar) L:M2 values for two different values of *k*_*transc*_ x two different values of *D*_*bias*_ under conditions of increased *P*_*transc*_ (= max of 0.9). As predicted, an increased *P*_*transc*_ resulted in a further decreased L:M2. Simulations with *D*_*bias*_ = 2 (results highlighted in pale yellow) were chosen for subsequent simulations. Right panel: histograms show simulated (blue bar) and experimentally observed (horizontal black bar) L:M2 values for two different values of *k*_*transc*_ x two different pol footprint sizes (14 and 7 nt) and *D*_*bias*_ = 2. A decreased pol footprint size increases the effective distance between the M2 GE and L GS signals, and results in simulated levels of L:M2 that closely match experimental observations. Each data point is the average of three 100,000 event simulations. **(C)** Global fits of the RSV gene expression data improve with the introduction of *D*_*bias*_ and reduced pol footprint size. Simulations of RSV transcription were performed at two different values of *k*_*transc*_ x two different values of *pol footprint* and *D*_*bias*_ = 2. Histograms of mRNA # for each RSV gene divided by the total mRNA # depict results from the simulations (blue bars) and average published experimentally observed values (black horizontal bars) (9). Each data point is the average of three 100,000 event simulations; error bars show the standard deviation. The number in parentheses and red above each histogram is the root-mean-square deviation (RMSD) of the simulated gene expression pattern from the experimental observations.

A 5’ pol diffusion bias was modeled by including a new parameter in the model, D_Bias_, with a value used as a multiplicative factor for 5’ diffusion only (Fig6A). Thus, a D_Bias_ value of 2 would result in a pol moving two steps (genomic chunks) with every 5’ movement while moving only one step (assuming D_scan_ = 1) with every 3’ movement; the probabilities of moving in either direction remain equal. This change could also be modeled by modifying the probabilities of 5’ vs. 3’ pol translocation and keeping each step size the same.

In order to test the effects of 5’ biased pol diffusion on gene expression in our model, we chose two of the parameter sets yielding fits with lower RMSDs from our first set of RSV transcription simulations involving multiple pols (Fig3B), ran these with three different values of D_Bias_, and looked for a drop in the predicted value of L:M2 mRNA levels (Fig6B). The two lower values of D_Bias_ tested produced a greater drop in L:M2 than the highest value tested (Fig6B). This is because under a maximum transcription initiation probability of 0.5, a high 5’ D_Bias_ leads to frequent ‘missing’ of the M2 GS signal before transcription initiation at the L GS signal. We therefore decided to increase the maximum transcription initiation probability to 0.9 and reran simulations at the lower values of 5’ D_Bias_ tested. As expected, this resulted in a further drop in predicted L:M2 mRNA levels. However, simulated L:M2 levels remained much higher than our experimentally observed value (Fig6B). Finally, we decided to decrease the pol footprint size by factors of 2 and 4, separately, knowing that this would increase the effective distance between the M2 GE signal and the L GS signal, and predicting a drop in L:M2 levels. The smallest pol footprint size tested, seven nts, is equal to the number of nucleotides bound by a single subunit of RSV nucleoprotein (N protein) and only three nts less than the size of the highly conserved RSV GS signal. Decreasing the pol footprint size yielded predicted L:M2 values that are very close to the experimentally observed value (Fig6B); and global fits of the RSV gene expression data quantitatively improved for the higher value of k_transc_ tested and remained roughly the same for the lower value (Fig6C).

### v. Optimizing model fits

With the addition of D_Bias_ to our model of RSV transcription, it seemed that both RSV and VSV versions of the model were poised to capture experimentally observed patterns of gene expression. We therefore set about finding RSV and VSV transcription initiation probabilities that would produce optimal fits of the experimental data (Fig7A and B). Using a set of transcription probabilities spanning a 10-fold range of values for a maximum pol number of 5, our RSV model yielded high quality fits of our experimental data (Fig7A and Table 2). Our VSV model yielded a high quality fit of the experimental data with a set of transcription probabilities spanning a 6-fold range and a maximum pol number of 50 (Fig7B and Table 2).

**Fig 7.**
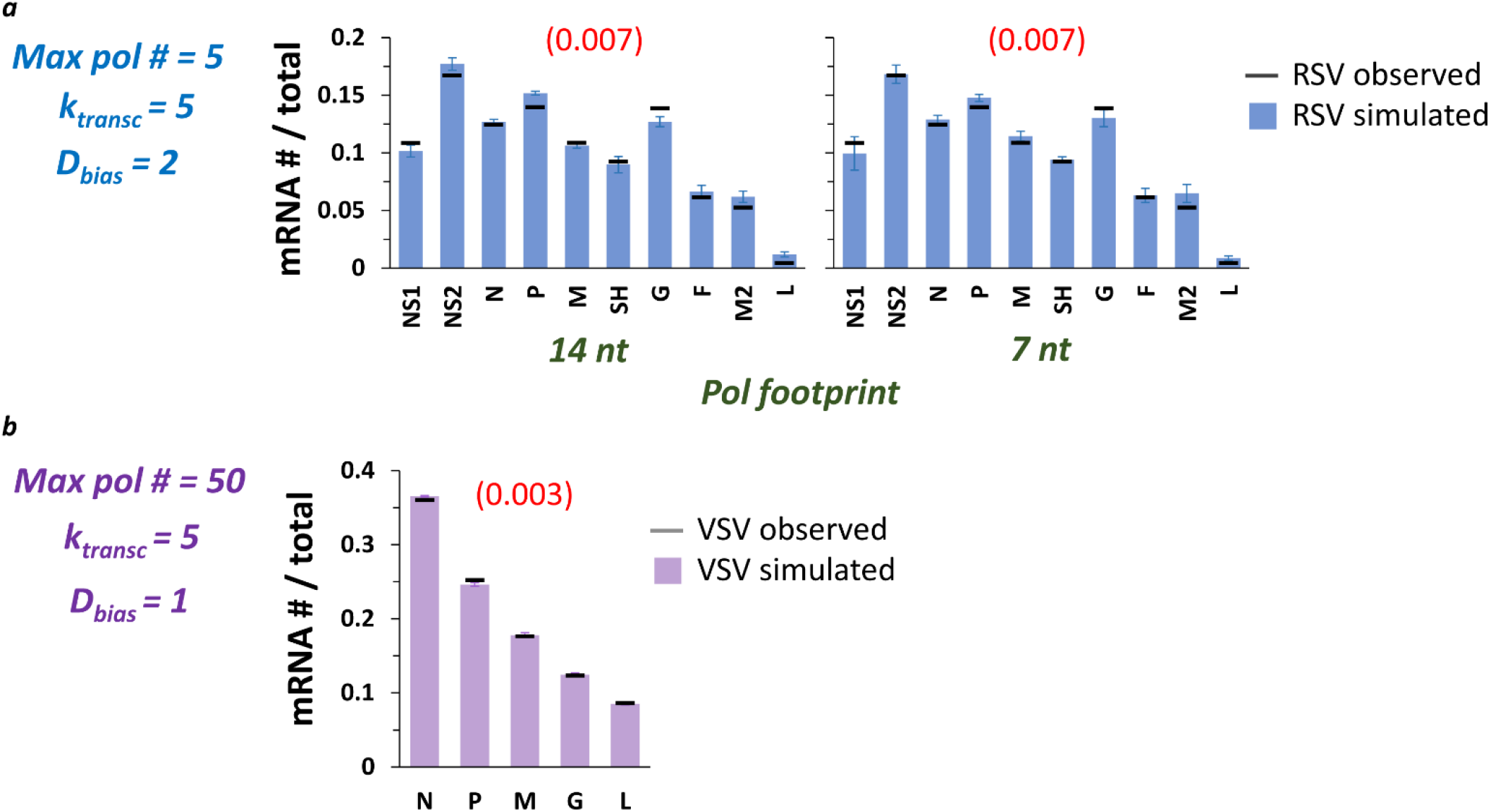
The model captures published observations of RSV and VSV transcription with adjustments to the underlying transcription probabilities (*P*_*transc*_). **(A)** High quality fits of experimentally observed RSV gene expression patterns. *P*_*transc*_ were manually adjusted to achieve optimized fits at *max pol #* = 5, *k*_*transc*_ = 5, *D*_*bias*_ = 2, and *pol footprint* of 14 and 7 nt. Histograms of mRNA # for each RSV gene divided by the total mRNA # depict results from the simulations (blue bars) and average published experimentally observed values (black horizontal bars) (9). Each data point is the average of three 100,000 event simulations; error bars show the standard deviation. The number in parentheses and red above each histogram is the root-mean-square deviation (RMSD) of the simulated gene expression pattern from the experimental observations. **(B)** A high quality fit of the benchmark experimentally observed VSV gene expression pattern. *P*_*transc*_ were manually adjusted to achieve an optimized fit at *max pol #* = 50, *k*_*transc*_ = 5, *D*_*bias*_ = 1 (i.e., NO 5’ bias), and *pol footprint* = 28 nt. The histogram of mRNA # for each VSV gene divided by the total mRNA # depicts results from the simulations (lavender bars) and average published experimentally observed values (black horizontal bars) (12). Each data point is the average of three 100,000 event simulations; error bars show the standard deviation. The number in parentheses and red above the histogram is the root-mean-square deviation (RMSD) of the simulated gene expression pattern from the experimental observations.

**Table 2.**
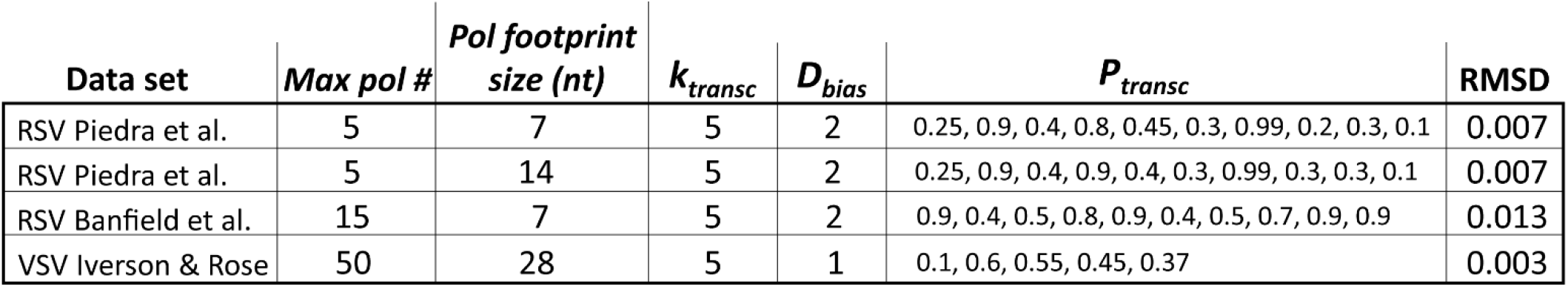
Major parameter values for high quality fits of different data sets by our model. Our model produces high quality fits of two different RSV data sets (4, 9) and benchmark observations of VSV gene expression (12). Each list of transcription initiation probabilities (*P*_*transc*_) contains the values used for every RSV or VSV GS signal following their 3’ to 5’ order along the genome.

We also decided to fit the recently reported RSV long-read sequencing data of Banfield et al (4). An increased *max pol #* and an approximately 2-fold range of *P*_*transc*_ were needed to capture their data (Table 2). These changes reflect the more gradient nature of the observed gene expression pattern, while our experimental observations showed much higher levels of G gene mRNA (9).

A 5’ diffusion bias was needed to capture both RSV data sets because of a common dramatic decrease in expression between genes M2 and L. In contrast, a 5’ diffusion bias was not needed to capture the benchmark observations of VSV gene expression used here; however, including one has minimal effect on the model’s output (data not shown). Thus, we simply cannot make a model-supported prediction about whether non-transcribing VSV pols diffuse with a 5’ bias. Continuing with VSV, the high quality fit we report involves a 6-fold range of *P*_*transc*_, but a quality fit can also be obtained with a 5-fold range of *P*_*transc*_ and less variation (= 0.1, 0.5, 0.5, 0.5, 0.5; RMSD = 0.009).

We believe the changes to transcription probabilities needed to produce high quality fits of the experimental data are reasonable. For instance, we have obtained preliminary data using RSV minigenomes encoding luciferase reporter genes showing that a single RSV GS signal sequence can support a 1.5-fold range of gene expression according to its alignment with bound nucleoprotein or *N-phase* (18). We do not know whether the reported N-phase-mediated changes to gene expression are exactly proportional to the changes in microscopic probabilities of transcription initiation modeled here because the former come from luciferase activity measurements and therefore reflect the addition of translation. Moreover, sequence changes outside of the highly conserved 10 nt stretch of the RSV GS signal can lead to gene expression changes (11), and the VSV GS signal is less conserved than RSV’s. However, we are not aware of minigenome studies exploring the effects of VSV GS signal sequence or N-phase on gene expression. Finally, it is also possible that the shape of the observed RSV and VSV gene expression patterns depends partly on differences in the underlying mRNA stabilities, which we make no attempt to model here; but we have shown previously that any such differences are unlikely to significantly affect experimentally observed RSV gene expression patterns (8).

### vi. Conclusions and limitations

Our model can capture observed RSV and VSV transcription patterns with biophysically reasonable parameters and parameter values. Our model makes the following major predictions in need of wet lab experimental testing: 1) ejective collisions occur between transcribing and non-transcribing NNSV pols; 2) non-transcribing RSV pols (and perhaps VSV pols) undergo 5’ biased diffusion along the viral genome; and 3) an increase in the number of pols bound to and diffusing along an NNSV genome at any one time will lead to more frequent pol-pol collisions and a sharper transcription gradient. Sophisticated single molecule TIRF-based assays are needed to directly test predictions 1-2, while 3 can be tested using established minigenome or recombinant genome assays along with high throughput sequencing.

## Methods

### The model

Computational models of RSV and VSV transcription were written in the Python programming language using the free and open-source Scientific Python Development Environment (Spyder version 3.3.2). The model code is freely available on GitHub: https://github.com/BCM-GCID/Publications/tree/main/Rethinking_NNSV_Gene_Expression.

In brief, the models simulate one or more viral RNA-dependent RNA polymerases (pols) entering a linear RSV or VSV genome at the 3’ end and taking a random walk at a rate D_scan_ (units = ‘genomic chunks’ per simulated event; D_scan_ = 1 throughout the results presented in this MS). A parameter D_bias_ is used as a multiplicative factor (D_scan_*D_bias_) to 5’ bias (or not) the random walk taken by modeled pols – i.e., D_bias_ > 1 biases pol movement 5’; D_bias_ =1 results in an unbiased random walk. Each genome is divided into chunks of a size thought to reasonably approximate the footprint of a single RSV or VSV pol (28, 14, or 7 nt). Diffusing non-transcribing pols cannot ‘hop’ over other pols and a single genomic chunk can only be occupied by a single pol at any one time.

Gene start (GS) and gene end (GE) signals are modeled as separate genomic chunks positioned along the modeled genomes according to their known positions from sequencing data (Fig1A). Transcription is initiated with a data-constrained probability (see Table 1) when a non-transcribing pol (pol_state = 0) diffuses onto a GS signal; termination of transcription or transcriptional readthrough occurs with a probability derived from published sequencing data when a transcribing pol (pol_state = 1) moving 5’ at a rate k_transc_ (units = ‘genomic chunks’ per simulated event) translocates onto a GE signal (Fig1B). Initiations of transcription and transcriptional readthrough events are counted as gene expression events for the genes where they occur. For simulations incorporating multiple pols on a single genome, ejections of a non-transcribing pol occur when a transcribing pol passes it. When a pol reaches the extreme 5’ end of a modeled genome, it either diffuses 3’ or dissociates from the genome.

The simulations occur one event at a time (i.e., time is modeled implicitly) whereby the positions and states (non-transcribing or transcribing) of the one or more modeled pols is stochastically updated according to the rules outlined above before proceeding to the next event. After simulating 10s of thousands of events, each gene’s mRNA level divided by the total mRNA level is outputted. These data are plotted to visualize a gene expression pattern.

## Acknowledgements

Thanks to Dr. Pedro A. Piedra and the Texas Medical Center Genomic Center for Infectious Diseases (TMC-GCID; NIH Grant # U19AI144297) for supporting this work. Thanks also to Vipin K. Menon of Baylor College of Medicine’s (BCM) Human Genome Sequencing Center (HGSC) and Sara J. Cregeen of BCM’s Alkek Center for Metagenomics and Microbiome Research (CMMR) for uploading the manuscript’s code to GitHub as one of multiple projects within the TMC-GCID.

